# Controlling complex microbial communities: a network-based approach

**DOI:** 10.1101/149765

**Authors:** Marco Tulio Angulo, Claude H. Moog, Yang-Yu Liu

## Abstract

Microbes comprise nearly half of all biomass on Earth. Almost every habitat on Earth is teeming with microbes, from hydrothermal vents to the human gastrointestinal tract. Those microbes form complex communities and play critical roles in maintaining the integrity of their environment or the well-being of their hosts. Controlling microbial communities can help us restore natural ecosystems and maintain healthy human microbiota. Yet, our ability to precisely manipulate microbial communities has been fundamentally impeded by the lack of a systematic framework to control them. Here we fill this gap by developing a control framework based on the new notion of structural accessibility. This framework allows identifying minimal sets of “driver species” through which we can achieve feasible control of the entire microbial community. We numerically validate our control framework on large microbial communities, and then we demonstrate its application for controlling the gut microbiota of gnotobiotic mice infected with *Clostridium difficile* and the core microbiota of the sea sponge *Ircinia oros*.

## INTRODUCTION

Microorganisms form complex communities that play critical roles in maintaining the well-being of their hosts or the integrity of their environment^1-4^. This deep relationship can have severe consequences to the host or the environment when a microbial community is disrupted. In humans, for example, a disruption to the gut microbiota —the aggregate of microorganisms residing in our intestine— has been associated to gastrointestinal diseases such as irritable bowel syndrome, and *Clostridium difficile* Infection (CDI)^5, 6^. A variety of non-gastrointestinal disorders as divergent as autism, obesity, and cavernous cerebral malformations have also been associated with disrupted gut microbiota^5, 7^. For agriculture crops, a disruption to rhizosphere microbiota can reduce their disease resistance and hence affect the overall crop yield^8, 9^. In the oceans, a disruption to their microbiota can impact global climate by altering carbon sequestration rates^3, 4, 10^. Driving these microbial communities back to their healthy states has the potential to bring novel solutions to prevent and treat complex human diseases, enhance sustainable agriculture, and regulate global warming^11, 12^. For example, inoculation of soil microbes can restore terrestrial ecosystems^13^, and Fecal Microbiota Transplantation (FMT) is so far the most successful therapy in treating patients with recurrent CDI by restoring disrupted gut microbiota^14^. Despite the success of these two empirical strategies, a broad application of microbial-manipulation strategies will only be possible if we can efficiently and systematically control large complex microbial communities^15^.

There are two main challenges to efficiently control a large complex microbial community. First and foremost, an efficient control method should only manipulate the minimal necessary number of species in the community. However, we still lack a method to systematically identify minimal sets of those “driver species” whose control can help us drive the whole community to desired states. Here, we use the term “species” in the general context of ecology, i.e., as a set of organisms adapted to a particular set of resources in the environment. It doesn’t necessarily represent the lowest major taxonomic rank. In fact, one could think of organizing microbes by strains, genera, or operational taxonomical units as well. Second, even if those driver species were known, calculating the control strategy that should be applied to them for driving the community towards the desired state remains somewhat tricky (e.g., it is difficult to calculate how much the abundance of those drive species needs to be increased or decreased). The difficulty in solving this second challenge is not only due to our insufficient knowledge of microbial dynamics and interactions, but also because of the inherently complex dynamics they often display.

To efficiently and systematically control large complex communities, here we develop a framework showing that the above two challenges can be addressed by focusing on the ecological network underlying the microbial community. We first introduce the new notion of “structural accessibility” and derive its graph-theoretical characterization. This theoretical result enables us to efficiently identify minimal sets of driver species of any microbial community purely from the topology of its underlying ecological network, even if some microbial interactions are missing and its population dynamics is unknown. Structural accessibility is a generalization of the notion of structural controllability^16^ —which only applies to systems with linear dynamics— to systems with nonlinear dynamics. Linear structural controllability is receiving increasing attention from the viewpoint of Network Science^17^. Once the driver species are identified, we systematically design feedback control strategies to drive a microbial community towards the desired state, even if the microbial dynamics is not precisely known. We numerically validated our control framework in large microbial communities, analyzing its performance for different parameters of the community we aim to control (e.g., the connectivity of its underlying ecological network), and with respect to errors in the ecological network used to identify the driver species. Finally, we demonstrate our framework by controlling the core microbiota of the sea sponge *Ircinia oros*, and restoring the gut microbiota of gnotobiotic mice infected by *Clostridium difficile*. Our results provide a rational and systematic framework to control microbial communities and other complex ecosystems based only on knowing their underlying ecological networks.

## PROBLEM STATEMENT

In our modeling framework, we focus on exploring the impact that manipulating a subset of species has on the abundances of other species. We thus consider a microbial community whose *state* at time *t* can be determined from the abundance profile *x*(*t*) ∈ ℝ^*N*^ of its *N* species, where the *i*-th entry *x_i_*(*t*) of *x*(*t*) represents the abundance of the *i*-th species at time *t*. Let us assume that the state evolves according to some general population dynamics

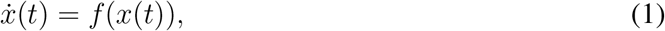

where the function *f*: ℝ^*N*^ → ℝ^*N*^ models the intrinsic growth and inter/intra-species interactions of the community (see Supplementary Note 1 for details). For most microbial communities the function *f* is unknown and difficult to infer due to the manifold of interaction mechanisms between microbes, such as cross-feeding and modulation by the host immune system^18^. Thus we assume that *f*(*x*) is some unknown *meromorphic* function (i.e., each entry *f_i_*(*x*) is the quotient of analytic functions of *x*). This is a very mild assumption that is satisfied by most population dynamics models^19^.

Instead of knowing the population dynamics of the microbial community, we assume we know its underlying *ecological network* 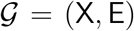. This network is defined as a directed graph where nodes X = {x_1_, ⋯, x_*N*_} represent species and edges (x_*j*_ → x_*i*_) ∈ E denote that the *j*-th species has a direct ecological impact (e.g., direct promotion or inhibition) on the *i*-th species (Fig.1a). Mapping these ecological networks requires performing mono-culture and co-culture experiments^20, 21^, using time-resolved abundance data and system identification techniques^22, 23^, or using steady-state abundance data via a recently developed inference method^24^. The accuracy of all these methods strongly depends on how informative is the available data^25^. Note that these ecological networks are different from correlation or co-occurrence based networks because correlation doesn’t imply causation^26^. Correlation-based networks can be readily constructed from abundance profiles of different samples^20, 27^ and, under certain specific conditions^28^, they could be a proxy of the underlying ecological network.

**Figure 1.**
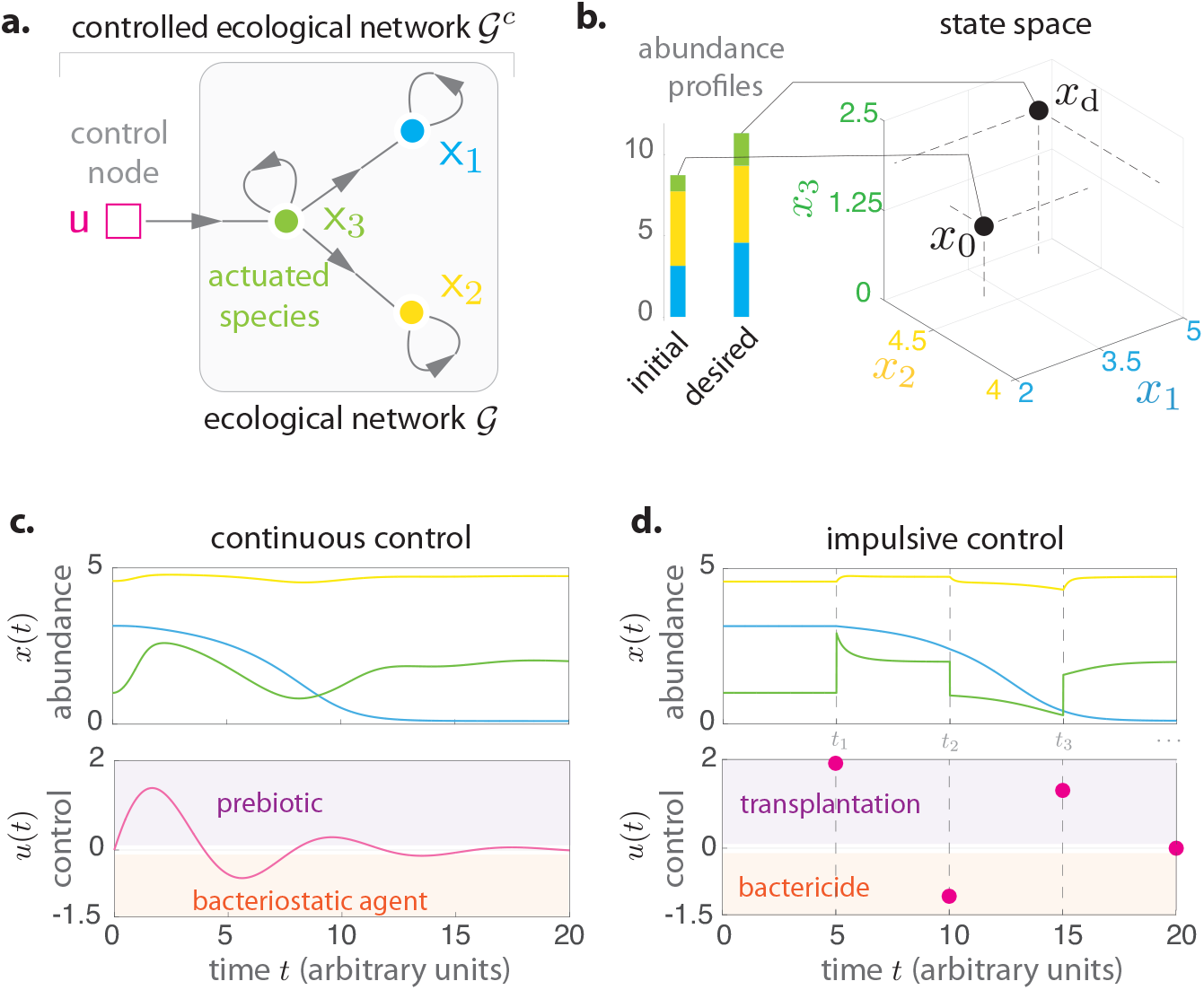
Controlling a microbial community. **a.** Ecological network 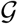 for a toy microbial community of *N* = 3 species (green, yellow, blue). The controlled ecological network 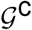 contains *M* = 1 control input actuating the third species. **b.** Initial and desired abundance profiles (bars). Controlling the community consists in driving its state from the initial state *x*_0_ to the desired state *x*_d_, represented by two points in the state space of the community. **c.** In the continuous control scheme, the control inputs *u*(*t*) are continuous signals modifying the growth of the actuated species. The controlled population dynamics of this community is given by 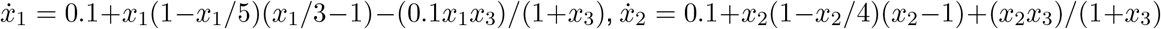, 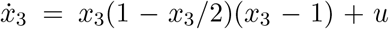. In the absence of control, this community has two equilibria *x*_0_ = (3.14, 4.58, 1)^T^ and *x_d_* = (4.57, 4.73, 2)^T^, chosen as the initial and desired states, respectively. **d.** In the impulsive control scheme, the control inputs *u*(*t*) are impulses applied at the intervention instants 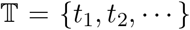, instantaneously changing the abundance of the actuated species. The controlled population dynamics is the same as in panel c, except that 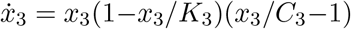 and 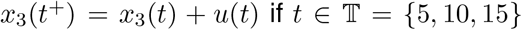. Under this controlled population dynamics, our mathematical formalism identifies x_3_ as the solo driver species needed to drive this microbial community (Example 1 in Supplementary Note 2).

Controlling a microbial community consists in driving its state from an initial value *x*_0_ = *x*(0) ∈ ℝ^*N*^ at time *t* = 0 (e.g., a “diseased” state) towards a desired value *x_d_* ∈ ℝ^*N*^ (e.g., a “healthier” state, Fig. 1b). We consider that the community will not naturally evolve to the desired state. To drive the microbial community, we consider a set of *M control inputs u*(*t*) ∈ ℝ^*M*^ that directly affect certain species that we call *actuated species* (Fig. 1a). These control inputs encode a combination of *M* control actions that are simultaneously applied to the community at time *t*. There are four types of control actions that we consider. If *u_j_* (*t*) < 0, the *j*-th control action at time *t* is either a *bacteriostatic agent* or a *bactericide*, which decreases the abundance of the species it actuates by inhibiting their reproduction or directly killing them, respectively^29^. If *u_j_* (*t*) > 0, the *j*-th control action at time *t* is either a *prebiotic*^30^ or a *transplantation*, which stimulate the growth or engrafts a consortium of the species it actuates, respectively. For the human gut microbiota, probiotics administration^31^ and FMTs^14^ are examples of transplantations. We introduce the *controlled ecological network* of the community 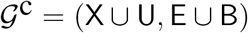 to specify which species are actuated by each control input. Here, the set U = {u_1_, ⋯, u_*M*_} is the set of control input nodes, and the edge (u_*j*_ → x_*i*_) ∈ B denotes that the the *j*-th control input actuates the *i*-th species (Fig.1a).

Given a controlled ecological network describing the interactions between species and which species are actuated by the control inputs, we next introduce two control schemes describing how the control inputs will affect the species. The first control scheme models a combination of prebi-otics (if *u_j_* (*t*) > 0) and bacteriostatic agents (if *u_j_* (*t*) < 0) as *continuous* control inputs modifying the growth of the actuated species (Fig. 1c):

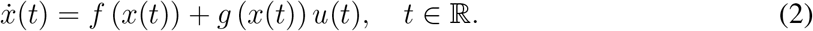

The second control scheme considers a combination of transplantations (if *u_j_* (*t*) > 0) and bactericides (if *u_j_*(*t*) < 0) applied at discrete *intervention instants* 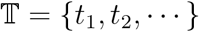, rendering *impulsive* control inputs that instantaneously modify the abundance of the actuated species (Fig. 1d):

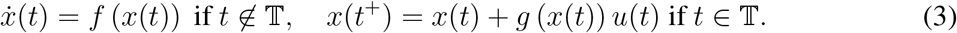

In the above equation, the symbol *x*(*t*^+^) denotes the state “right after time *t*”, so a control input *u*(*t*) ≠ 0 at 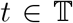 makes *x*(*t*) “jump” at that time instant. Thus, control actions are classified as impulsive if they instantaneously modify the abundance of some species, and continuous otherwise (see Supplementary Note 1.2 for details).

Both control schemes are characterized by the pair of functions {*f*, *g*}, describing the controlled population dynamics of the microbial community. As we have seen, the function *f*: ℝ^*N*^ → ℝ^*N*^ models the intrinsic growth and inter/intra-species interactions. The function *g*: ℝ^*N*^ → ℝ^*N*×*M*^ models the direct susceptibility of the species to the control actions. The *i*-th species is actuated by the *j*-th control input if the (*i*, *j*)-th entry of *g*(*x*) satisfies *g_ij_*(*x*) ≢ 0. As in the uncontrolled community of Eq. (1), the function *g*(*x*) is typically unknown because the mechanisms of susceptibility to the control actions can be uncertain. Thus we assume that *g* is some unknown meromorphic function such that *g_ij_* ≢ 0 iff (u_*j*_ → x_*i*_) ∈ B.

Notice that when all species are directly controlled (i.e., each species is actuated by an independent control input so *M* = *N* and *g*(*x*) is full rank), the state of the whole microbial community can obviously be fully controlled. Fortunately, as we next show, controlling all the species in a community is far from being necessary. Indeed, several species can be *indirectly* controlled by the same control input when this signal is adequately propagated through the ecological network underlying the community. Thus, our first goal is to identify minimal sets of species that we need to actuate in order to drive the entire community. We call those species *driver species*. We will also study if the impulsive control scheme can be as effective as the continuous control scheme for controlling microbial communities. Indeed, the former is more feasible than the latter, especially for human-associated microbial communities. Finally, we will design the control inputs that should be applied to the identified driver species to drive the whole community towards the desired state.

## IDENTIFYING DRIVER SPECIES

### Driver species are characterized by the absence of autonomous elements

To understand when a set of actuated species is a set of driver species, consider a three-species community with the classical Generalized Lotka-Volterra (GLV) population dynamics (Fig. 2a). This toy community has one control input actuating the third species x_3_. Actuating this species alone creates an *autonomous element* —namely, a constraint between some species abundances that the control input cannot break, confining the state of the community to a low-dimensional manifold. More precisely, our mathematical formalism reveals that ξ = *x*_1_ *x*_2_ is an autonomous element for this microbial community (Example 2 in Supplementary Note 2). Indeed, differentiating ξ with respect to time yields 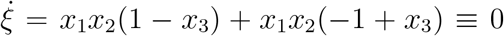, which implies that the state of the community is constrained to the low-dimensional manifold {*x* ∈ ℝ^3^|*x*_1_*x*_2_ = *x*_1_(0)*x*_2_(0)} for all control inputs (Fig. 2a right). Intuitively, an autonomous element exists because the control input cannot change the abundance of species x_1_ without changing the abundance of species x_2_ in a predefined way (i.e., *x*_2_ = *x*_1_(0)*x*_2_(0)/*x*_1_). It is thus impossible to drive the whole community in its three-dimensional state space, implying that x_3_ cannot be a driver species for this community. Introducing a second control input actuating species x_1_ eliminates this autonomous element by helping the system to jump out of the low-dimensional manifold. Hence, the community can be driven in any direction within its three-dimensional state space (Fig. 2b). This indicates that {x_1_, x_3_} is a minimal set of driver species for this community. Actually, by using these two driver species we can steer the community to any desired state with positive abundances (Example 6 in Supplementary Note 5).

**Figure 2.**
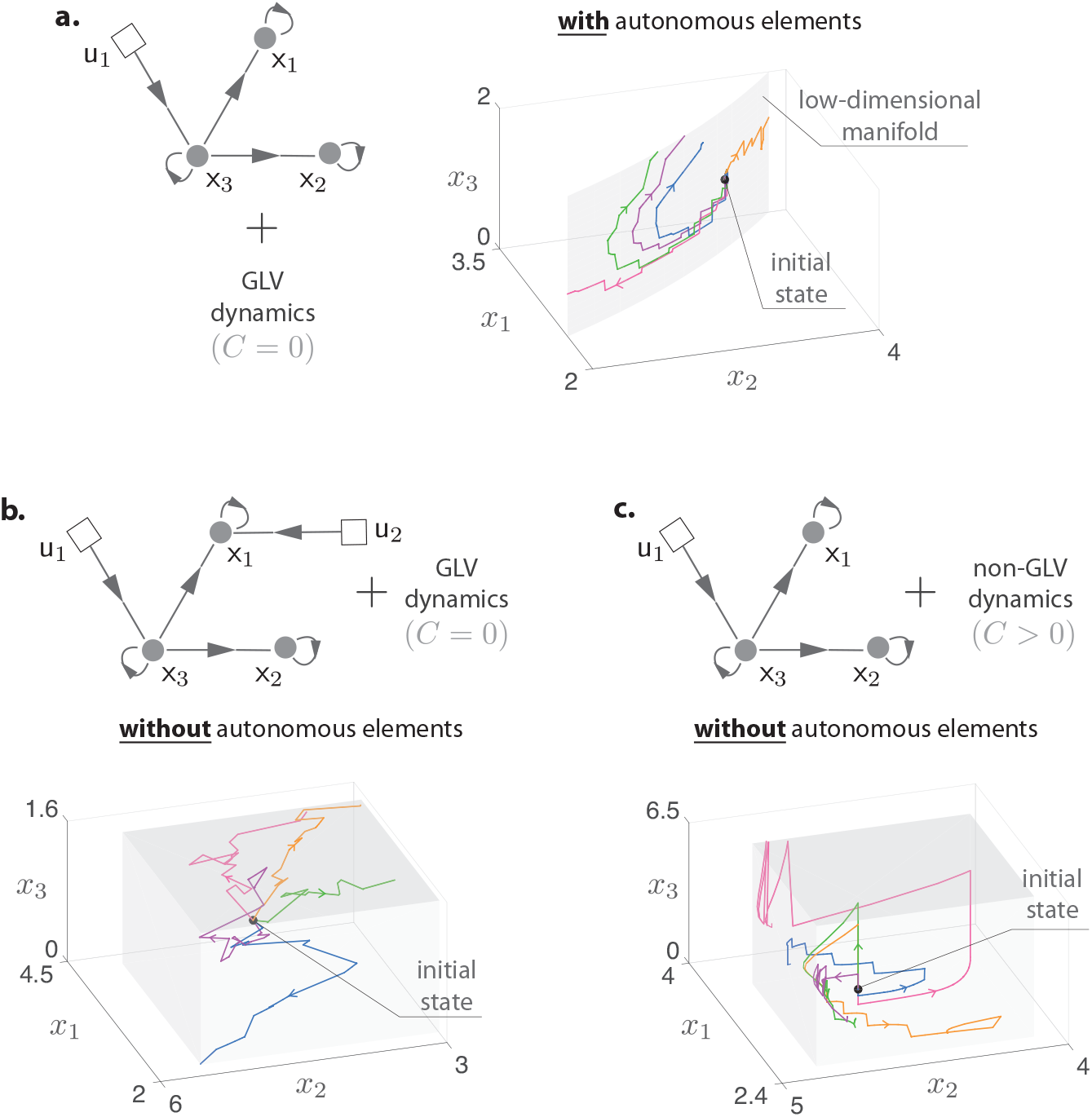
Autonomous elements constrain the state of microbial communities, characterizing their driver species. **a.** A three-species community with GLV dynamics 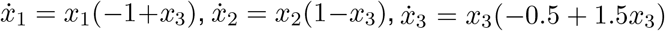. For actuating x_3_, we consider the impulsive control scheme with *x*_3_(*t*^+^) = *x*_3_(*t*) + *u*_1_ (*t*) for 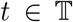. With this controlled population dynamics, our mathematical formalism reveals the autonomous element *x*_1_*x*_2_ that constraints the state of this microbial community to the low-dimensional manifold {*x* ∈ ℝ^3^|*x*_1_*x*_2_ = *x*_1_(0)*x*_2_(0)} (gray) for all control inputs. Five state trajectories (in colors) with random control inputs illustrate this fact. Hence, {x_3_} alone cannot be a set of driver species for this controlled population dynamics. **b.** Including a second control input *u*_2_(*t*) actuating x_1_ (i.e., *x*_1_ (*t*^+^) = *x*_1_(*t*) + *u*_2_(*t*) for 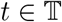) eliminates the autonomous element, since the state of the microbial community (colors) can explore a three-dimensional space (gray). Hence {x_1_, x_3_} is a minimal set of driver species for this community with GLV dynamics. **c.** We proved that, generically, increasing the complexity of the controlled population dynamics cannot create autonomous elements. In this example, increasing the deformation size *C* from the GLV in panel a (with *C* = 0) to the controlled population dynamics in Fig.1 (with *C* > 0) eliminates the autonomous element that was present by actuating x_3_ alone (Example 1 in Supplementary Note 2). Therefore, increasing the complexity of the population dynamics makes {x_3_} a solo driver species.

In the general case of *N* species and *M* control inputs, we define a set of actuated species as a set of driver species if the corresponding controlled population dynamics {*f*, *g*} of the microbial community lacks autonomous elements. For a given pair {*f*, *g*}, the absence of autonomous elements can be mathematically deduced using a formalism based on differential one-forms (Supplementary Note 2). Indeed, for the continuous control scheme of Eq. (2), the conditions for the absence of autonomous elements are well understood because they define when a system is *accessible*^32^. As a cornerstone concept in nonlinear control theory, accessibility has been instrumental for developing technological advances such as robotics. Since it is more natural to control microbial communities with impulsive control actions, in this paper we extended the study of autonomous elements to the impulsive control systems of Eq. (3). For this, we first introduced a mathematical definition of autonomous elements for impulsive control systems (Definition 3 in Supplementary Note 2). Using this definition, we characterized necessary and sufficient conditions for the absence of autonomous elements in a given controlled population dynamics (Theorem 2 in Supplementary Note 2).

To our surprise, we found that the conditions for the absence of autonomous elements for the continuous and the impulsive control schemes are identical (Remark 2 in Supplementary Note 2). This result suggests that, for controlling microbial communities, transplantations and bactericides (impulsive control actions) can be as effective as prebiotics and bacteriostatic agents (continuous control actions). Since impulsive control actions could be simpler to implement for many microbial communities such as the human gut microbiota, this result assures us to further develop microbiome-based therapies in the form of probiotic cocktails and FMTs.

### Structural accessibility characterizes the generic absence of autonomous elements

For complex microbial communities such as the human gut microbiota, it is very difficult to choose an adequate pair {*f*, *g*} to model its controlled population dynamics. As the autonomous elements depend on such a pair, this might suggest that it is impossible to predict their presence and thus to identify the driver species of complex microbial communities. We now show that this seemingly unavoidable limitation can be solved by focusing on the topology of the controlled ecological network of the community.

Define the graph 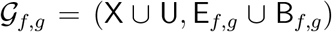 associated with a meromorphic function pair {*f*, *g*} as follows. First, the edge (x_*j*_ → x_*i*_) ∈ E_*f*,*g*_ exists if *x_j_* appears in the right-hand side of 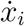 or *x_i_*(*t*^+^) in Eqs. (2) or (3), respectively. Second, the edge (u_*j*_ → x_*i*_) ∈ B_*f*,*g*_ exists if *g_ij_* ≢ 0. In this definition, the interaction x_*j*_ → x_*i*_ can originate in the uncontrolled population dynamics (i.e., *f_i_*(*x*) depends on *x_j_*) or, in a more general case, also in the controlled dynamics (i.e., the *i*-th row of *g*(*x*) depends on *x_j_*). Using this definition and given a controlled ecological network 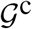, we can describe the class 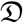 of all possible controlled population dynamics that the controlled microbial community can have. Mathematically, we describe the class 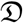 as containing all *base models* {*f*\*, *g*\*} such that 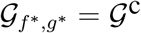, together with all *deformations* {*f*, *g*} of each of those base models. The base models characterize the simplest controlled population dynamics that the community can have. We have chosen them as controlled GLV models with constant susceptibilities:

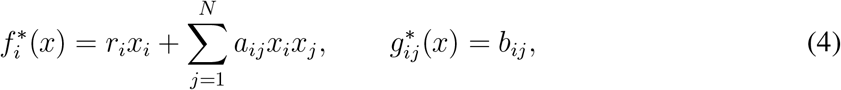

for *i* = 1, ⋯, *N*. The base models are parametrized by *A* = (*a_ij_*) ∈ ℝ^*N*×*N*^, *r* = (*r_i_*) ∈ ℝ^*N*^, and *B* = (*b_ij_*) ∈ ℝ^*N*×*M*^, representing the interaction matrix, the intrinsic growth rate vector, and the susceptibility matrix of the community, respectively. Thus, the base models in 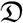 are all controlled GLV models such that their graph matches 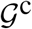. As a classical population dynamics model, the GLV model has been applied to microbial communities in lakes, soils, and human bodies^14, 15, 20, 33–39^. Notice that in a microbial community, any species that gets extinct cannot “resurrect” by itself without some external influence such as a transplantation or migration. Eq. (4) is the simplest population dynamics that satisfies this condition in the following sense: it is obtained by considering population dynamics of the form *f_i_*(*x*) = *x_i_F_i_*(*x*), and then choosing the functions *F_i_*(*x*) to be simple affine functions.

Next, we say that a meromorphic pair {*f*, *g*} is a deformation of a base model {*f*\*, *g*\*} if it satisfies the following three conditions: (i) it has the same graph as the base model (i.e., 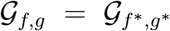); (ii) there exists a finite set of parameters *θ* ∈ ℝ^*C*^ such that 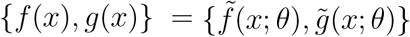; and (iii) the identity 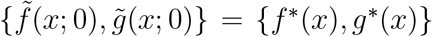 holds. The minimal integer *C* ≥ 0 for which these conditions are satisfied is called the *size* of the deformation, quantifying the cardinality of the parameter set *θ* that is needed to obtain the deformation from the base model. A rather general class of controlled population dynamics can be described by deformations of the base model of Eq. (4), such as

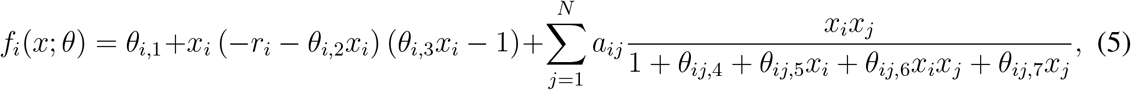

for *i* = 1, ⋯, *N*. In Eq. (5), the parameters *θ*_*i*,1_ are migration rates from/to neighboring habitats, 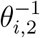 are the carrying capacities of the environment, 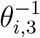 are the Allee constants, and the rest 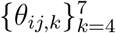 characterize the saturation of the functional responses^40^. Note that *θ*_*i*,1_ > 0 can also model species like *C. difficile* that sporulate into “inactive” forms and then recover. Note also that “higher-order” interactions can be described as deformations. For example, if species x_*i*_ is directly affected by species x_*j*_ and x_*k*_, then a deformation can include the third-order interaction *θ_i_x_i_x_j_x_k_*. Similarly, deformations allow cases when the susceptibility of the *i*-th species to *j*-th control input is mediated by the abundance of other species. For example, the deformation *g_ij_*(*x*; *θ*) = *b_ij_* + *θ_jik_x_k_* models a case when the *i*-th species is actuated by the *j*-th control input but its effect is mediated by the abundance of the *k*-th species.

We call the class 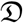 *structurally accessible* if almost all of its base models and almost all of their deformations lack autonomous elements. This means that, except for a zero-measure set of “singularities”, all the controlled population dynamics that the community may take have to lack autonomous elements. The conditions under which 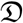 is structurally accessible are fully characterized using our mathematical formalism (Supplementary Note 3), and they depend only on the underlying controlled ecological network 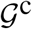. We first proved that, generically, increasing the size of a deformation cannot create autonomous elements (see Proposition 1 in Supplementary Note 3, and Fig. 2c for an illustration). This result reduces the search for autonomous elements to the deformations in 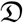 with minimal size *C* = 0. That is, to all base models whose graph matches 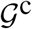. Finally, we proved that 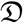 is structurally accessible if and only if 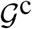 satisfies the following two conditions: (i) each species is the end-node of a path that starts at a control input node; and (ii) there is a disjoint union of cycles (excluding self-loops) and paths that cover all species nodes (see Theorem 3 of Supplementary Note 3). If these two graph conditions are satisfied, we also call 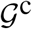 structurally accessible.

The notion of structural accessibility introduced above is a nonlinear counterpart of the notion of structural controllability for linear systems^16^. For linear systems we have {*f*(*x*), *g*(*x*)} = {*Ax*, *B*}, and the absence of autonomous elements is equivalent to their controllability^32^ —the intrinsic ability to drive the system between two arbitrary states, which can be verified by the celebrated Kalman’s rank condition: rank(*B*, *AB*, *A*^2^*B*,…, *A*^*N*−1^*B*) = *N*. Condition (i) above is necessary for both structural accessibility and linear structural controllability, requiring that the network contains paths that spread the influence of the control inputs to all species. However, for linear structural controllability, condition (ii) is sufficient but not necessary. More precisely, for linear structural controllability, the required disjoint union of cycles that cover the species nodes can also include self-loops due to intrinsic nodal dynamics (see Remark 4 in Supplementary Note 3).

### Identifying minimal sets of driver species in microbial communities

The above result provides a complete graph-characterization of driver species: a set of actuated species is a set of driver species (for all but a zero-measure set of controlled population dynamics that the community may have) if and only if its corresponding 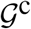 is structurally accessible. We used this characterization to build an algorithm that identifies a minimal set of driver species from the ecological network of the community. More precisely, we mapped the satisfaction of the graph conditions (i) and (ii) into solving a maximum matching problem over the graph 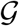 without self-loops (Proposition 3 in Supplementary Note 4). This result provides a polynomial time algorithm to identify one minimal set of driver species, making it feasible for large networks (Remark 5 in Supplementary Note 4).

Note that once 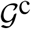 is structurally accessible this network cannot lose its structural accessibility when new edges are added to it. This observation implies that a set of driver species remains valid even if new edges (e.g., new inter/intra-species interactions) are added to the ecological network of the community. Therefore, it is possible to find the driver species of a microbial community using an “incomplete” ecological network that only includes some of the ecological interactions (e.g., high-confidence interactions).

## DRIVING THE DRIVER SPECIES

Next we turn to the question of calculating the control signal *u*(*t*) that needs to be applied to a set of driver species to drive the whole community towards the desired state. We will show that impulsive control actions can make this calculation easier.

### Calculating optimal control strategies for microbial communities with known population dynamics

To calculate the impulsive control inputs 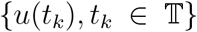 needed to drive the microbial community to the desired state *x_d_* we adopt a *model predictive control* (MPC) approach^41^. First, based on the current state of the community *x*(*t_k_*) at the intervention instant 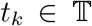, we use knowledge of its controlled population dynamics to predict the sequence of states 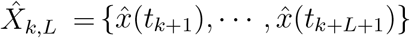 that the community will take in response to a sequence of *L* impulsive control inputs *U_k, L_* = {*u*(*t_k_*), ⋯, *u*(*t*_*k*+*L*−1_)}. The *prediction horizon L* > 0 quantifies how far into the future we predict. We then choose 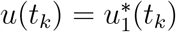, where 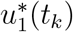 is the first element of the optimal control input sequence 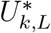 calculated by solving the following optimization problem:

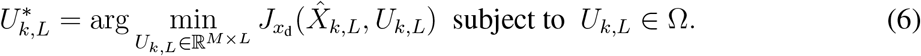

Here Ω ⊆ ℝ^*M*×*L*^ is a set that specifies constraints in the control inputs we can use, and *J_xd_* is some cost function penalizing deviations of the predicted trajectory 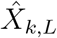 from the desired state *x_d_*. For example, the simplest cost function 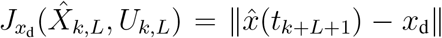 penalizes deviations of the predicted final state from the desired state. Penalizing the deviations of intermediate states can provide a smoother transition to the desired state.

To choose the prediction horizon *L* in Eq. (6), we proved that it is possible to distinguish between two cases (Theorem 4 in Supplementary Note 5). The first case is when the community can be driven to the desired state using a finite number *L* of impulsive control actions. This number can be calculated from its controlled population dynamics. The second case is when the community can only be asymptotically driven to the desired state as time goes by, meaning that a “sufficiently large” *L* ≫ *N* should be used. This second case could be circumvented by increasing the number of driver species (Remark 8 in Supplementary Note 5). Note that by recalculating 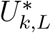 at each intervention instant 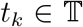 using the actual state of the community, the MPC method creates a feedback loop that enhances its robustness against prediction errors due to uncertainty in the dynamics^41^. For *L* = 1 the proposed MPC methodology is similar to the network control method of Ref. (40). Eq. (6) is a finite-dimensional optimization problem that can be solved using several algorithms such as “DIRECT”^42^. By contrast, for continuous control actions, the analogous optimization problem is defined over the infinite-dimensional space of all *M*-dimensional continuous functions. Solving such optimization problem is apparently more difficult, significantly limiting our ability to calculate optimal continuous control actions.

We studied the performance of the above MPC strategy in the three-species microbial community with a solo driver species of Fig. 1. Given the dynamics of this community (see caption in Fig. 1), we find that *L* = 3 impulsive control inputs are sufficient to drive the whole community (Example 4 in Supplementary Note 5). To calculate the optimal control inputs we selected 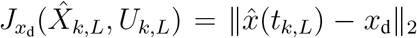 in Eq. (6). Solving the optimization problem using DIRECT yields the MPC strategy *u*\*(*t*_1_) = −0.8815, *u*\*(*t*_2_) = 2.0089 and *u*\*(*t*_3_) = −10^−4^ (pink in Fig. 3a). We use this example to compare the performance of applying two other control strategies to drive this community. The first strategy uses a transplantation to restore the abundance of the driver species (i.e., increase its abundance to its desired value), expecting that such control action will drive the rest of the community to the desired state (purple in Fig. 3a). This control strategy is reminiscent of a probiotic administration that restores the “healthy” abundance of the driver species. The second control strategy ignores the driver species of this community, using two control inputs (instead of one) to set the abundance of the non-driver species to their desired values (blue in Fig. 3a).

**Figure 3.**
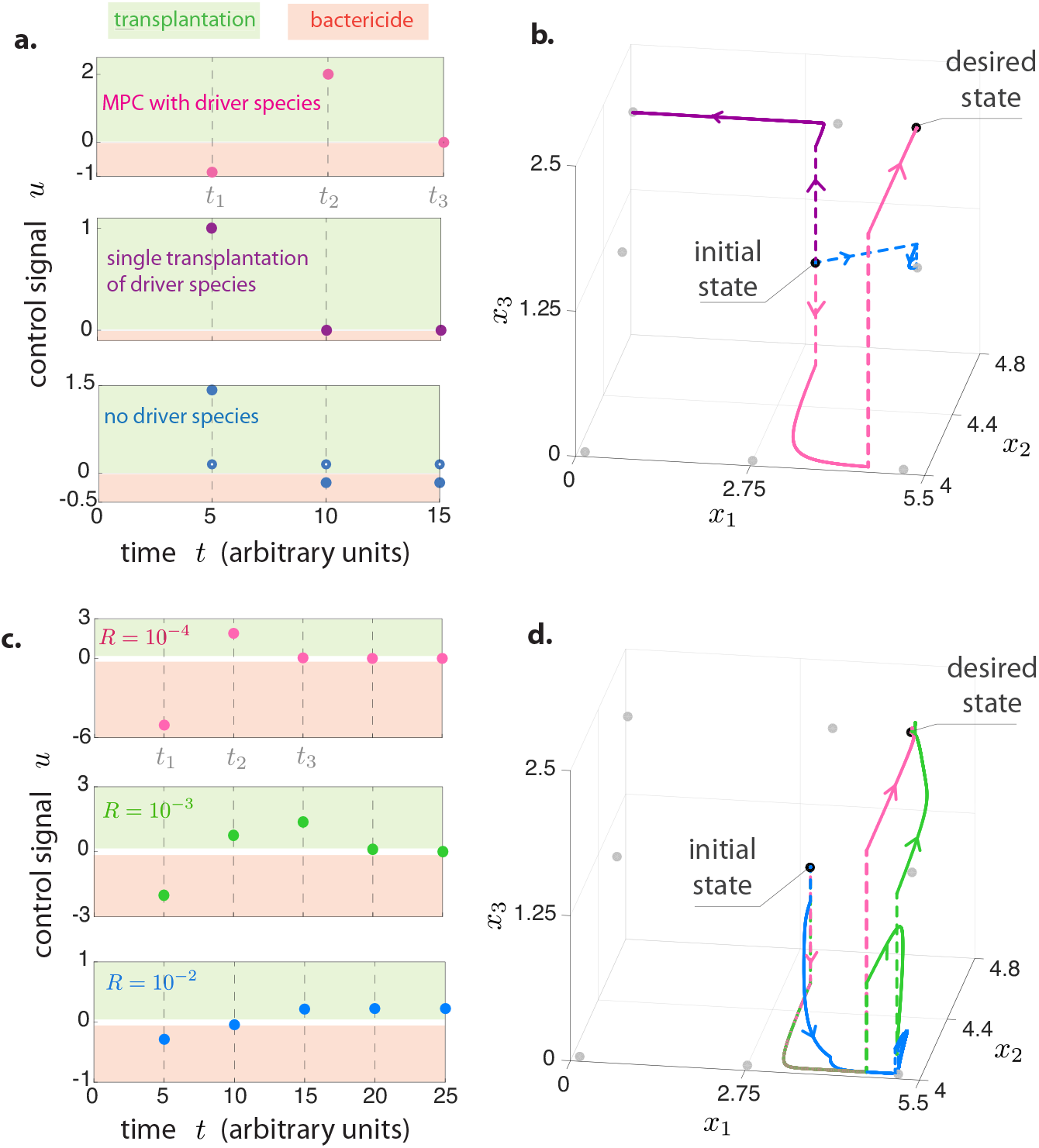
Success and failure of different control strategies. **a.** Three control strategies for driving the microbial community of Fig. 1a toward the desired state. First, MPC applied to the identified driver species {x_3_} (pink dots). The second control strategy increases the abundance of the driver species to match its value at the desired state *x*_3_(*t*_1_) = *x*_3,d_ (purple dots). The third control strategy does not actuate the driver species, but actuates the other two species {x_1_, x_2_} by setting their abundance to their desired values (i.e., *x*_1_(*t_k_*) = *x*_1, d_ and *x*_2_(*t*_k_) = *x*_2,d_, solid and hollow blue dots, respectively). **b.** The response of the microbial community to these three control strategies. Here and in panel d, the “jumps” produced by the control inputs are depicted by dashed lines. The equilibria of the population dynamics are shown as gray dots. Only the first strategy applying MPC to the driver species succeeds in driving the community to *x*_d_. **c.** Control strategies obtained by using the linear MPC with parameters *Q* = diag(20, 1, 10) and different values for *R*: 10 ^−4^ (pink), 10^−3^ (green), 10^−2^ (blue). **d.** Trajectories of the controlled community using the linear MPC control strategies described in panel c. Colors correspond to the different values of *R*.

Among the above control strategies, only the MPC applied to the driver species succeeds (Fig. 3b). Actually, this strategy succeeds in a somewhat unconventional way: despite the driver species is more abundant in the desired state than in the initial state, the first control action decreases further its abundance. This first control action makes the non-driver species reach their desired abundances and, once that happens, the abundance of the driver species is finally increased to its desired value (pink in Fig. 3b). The second control strategy succeeds in driving species x_2_ and x_3_, but it fails to drive x_1_ to the desired abundance because it approaches the desired state from an unstable direction (purple in Fig. 3b). Finally, not actuating the driver species results in the worst strategy, failing to drive a single species to the desired state (blue in Fig. 3b). This example demonstrates the importance of actuating the driver species.

### Calculating control strategies for microbial communities with uncertain population dynamics or a large number of species

In general, solving the non-convex optimization problem of Eq. (6) is challenging as the number of species or prediction horizon increase. Also, a prerequisite for solving this optimization problem is a reasonable knowledge of the controlled population dynamics of the community, which may not available. To circumvent these two drawbacks, next we leverage the network underlying the controlled microbial community.

Consider that it is possible to obtain a weighted adjacency matrix *Â* ∈ ℝ^*N*×*N*^ from the ecological network 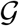 of the community, providing a proxy for its interaction matrix. Without additional knowledge of the susceptibility matrix of the community, we assume it is possible to increase or decrease as desired the abundance of each driver species. Under this assumption, we define 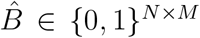 as a proxy for the susceptibility matrix, with *b_ij_* = 1 if the *j*-th control input actuates the *i*-th driver species. Next, by rewriting the controlled population dynamics of the community as 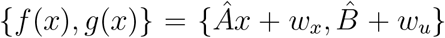, we use the pair 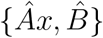 to provide a linear prediction for the response of the community to the control inputs. Here, the nonlinear functions 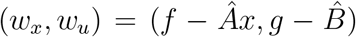 represent perturbations whose magnitude depend on how well the linear pair 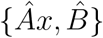 approximates the true dynamics {*f*(*x*),*g*(*x*)} of the community. Using this linear pair for predicting the response of the community to impulsive control actions, we design a *linear MPC* by solving the optimization problem of Eq. (6) with the quadratic cost function

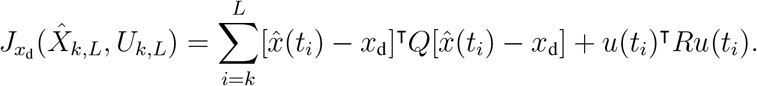

In the above equation, the positive definite matrices 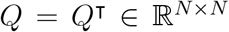 and *R* = *R*^T^ ∈ ℝ^*M*×*M*^ are design parameters. The matrix *Q* penalizes the deviations of the predicted trajectory from the desired state, and *R* quantifies the “cost” of using the control inputs. Under this scenario (i.e., a linear prediction model and quadratic cost), the solution to the optimization problem of Eq. (6) can be obtained in closed form^43^ even if *L* → ∞. This result enabled us to obtain the explicit form *u*(*t_k_*) = *Kx*(*t_k_*) for the linear MPC at time 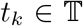, where *K* ∈ ℝ^*M*×*N*^ is computed by solving a Riccati algebraic equation (Supplementary Note 6). Since the Riccati equation can be efficiently solved for large *N*, the linear MPC can be calculated for large microbial communities. The above linear MPC has several other advantages: it requires minimal knowledge of the controlled population dynamics of the community (i.e., the weighted adjacency matrix of its underlying ecological network); it is robust to the perturbations (*w_x_*,*w_u_*) and other uncertainties (Remark 12 in Supplementary Note 6); and it also allows calculating the control signals for the continuous control scheme (Remark 10 in Supplementary Note 6).

We used the above linear MPC for controlling the three-species community of Fig. 1, assuming its dynamics is unknown. Based on the ecological network of this community and its population dynamics (see Fig. 1 and its caption), we choose *Â* = (−0.5,0, −0.1; 0, −5, 1; 0, 0, −1) as a proxy for its interaction matrix. Note that *Â* is a rather rough approximation of the linearization of the population dynamics at the desired state given by (−0.37, 0, −0.05; 0, −5.31, 0.52; 0, 0, −1). Since {x_3_} is a solo driver species for this community, we use 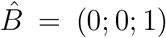. Choosing *Q* = diag(20, 1, 10), we compared the performance of three different linear MPCs obtained by using the values *R* = 10^−4^, 10^−3^, 10^−2^ (Fig. 3c). The performance of the linear MPC strongly depends on the selection of these parameters. For *R* = 10^−4^, despite not using knowledge of the population dynamics, the performance of the linear MPC (pink in Fig. 3d) is very similar to the performance of the MPC that uses full knowledge of the nonlinear population dynamics (pink in Fig. 3b). The success of the linear MPC in driving a community with nonlinear population dynamics illustrates the robustness of the MPC strategy, since the controller succeeds despite having non-zero perturbations (*w_x_*, *w_u_*). As *R* increases, the performance of the linear MPC deteriorates, first using more interventions to reach the desired state (green in Fig. 3d), and finally failing to drive the system to the desired state (blue in Fig. 3d). Indeed, since *R* > 0 quantifies the “cost” of using control inputs, increasing *R* reduces the magnitude of the control inputs, to the point they are not large enough to drive the system towards the desired state. We emphasize that, in general, the performance of the linear MPC also depends on the chosen 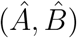 and the desired state (Remark 11 in Supplementary Note 6).

### Numerical validation of the control framework on large microbial communities

To systematically validate our control framework, we considered communities of *N* = 100 species having random Erdös-Rényi ecological networks with a prescribed connectivity *c* ∈ [0, 1], see Fig. 4a. The network edge-weights are chosen from a normal distribution with zero mean and standard deviation *σ* ≥ 0, where *σ* characterizes the typical interspecies interaction strength. Negative self-loops with weights −1 were added to each species to ensure stability, representing intraspecies interactions. We use this ecological network to identify the driver species of the community, and its corresponding weighted adjacency matrix as the interaction matrix to construct the linear MPC. The parameters *Q* = 20 × 10^4^*I*_*N*×*N*_, *R* = 0.15*I*_*M*×*M*_ of the linear MPC were fixed for all communities, and the intervention time instants 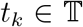 were chosen such that *t*_*k*+1_ − *t_k_* = 0.1. Next, we used Eq. (5) to numerically simulate the population dynamics of these communities. For this, we set the weighted adjacency matrix of the ecological network we built as the interaction matrix *A* in Eq. (5). We choose *θ*_i,j_ · = 0 for *j* = 1, ⋯, 6, and *θ*_*ij*,7_ uniformly at random from [0, *θ*_max_], where *θ*_max_ is a parameter. Last, we choose the intrinsic growth rates *r_i_* to ensure all generated random communities share the desired state *x*_d_ ∈ ℝ^*N*^ as an equilibrium point. Note all the constructed communities have nonlinear population dynamics, and their linearization at the desired state is not equal to the interaction matrix used for the linear MPC (see Supplementary Note 8 for details of this construction).

**Figure 4.**
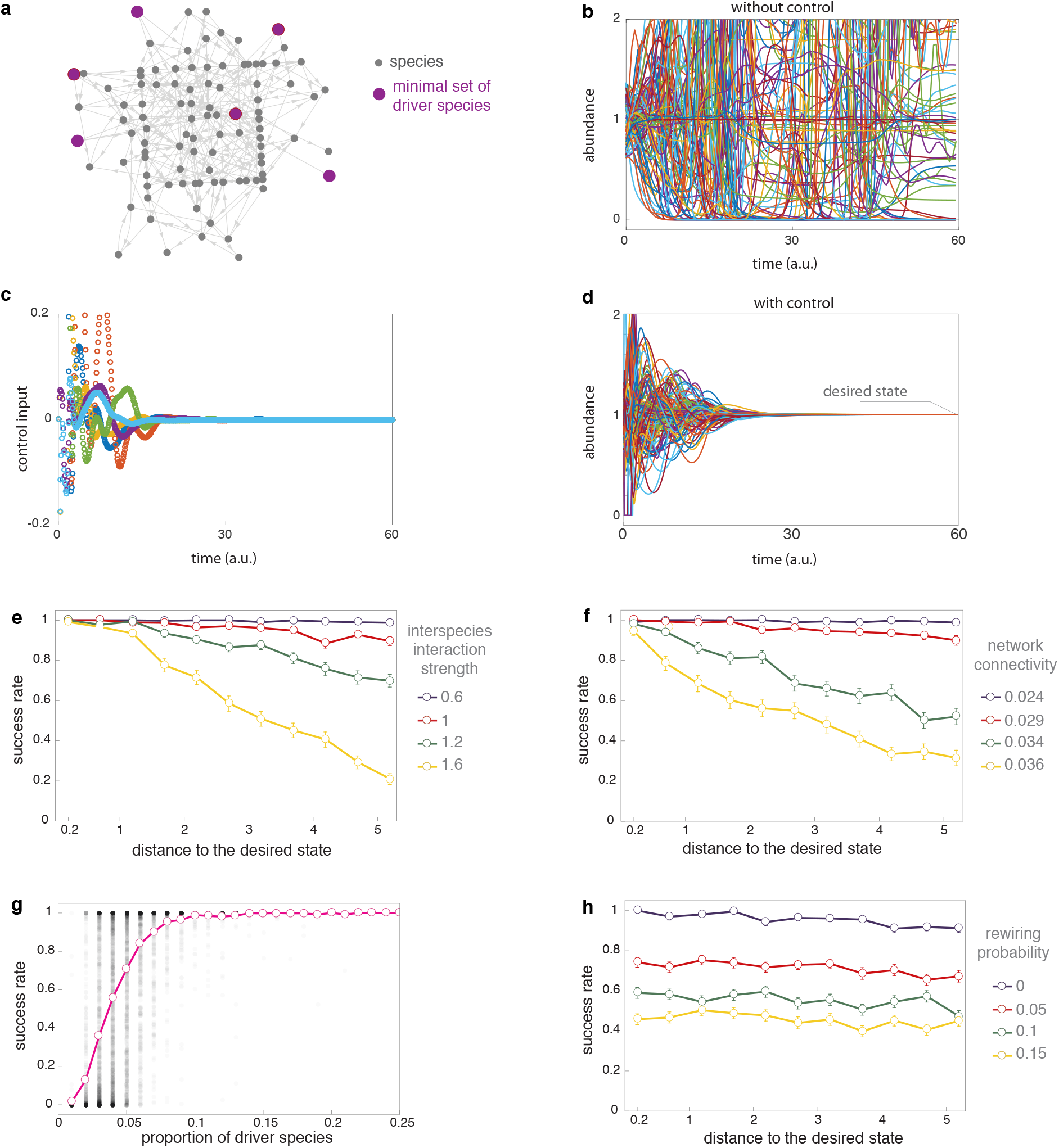
Numerical validation of the control framework on large microbial communities. **a.** Example of the ecological network of a random microbial community with *N* = 100 species with connectivity *c* = 0.03. We used our framework to identify a minimal set of *M* = 6 driver species. The desired state is chosen as *x*_d_ = (1, ⋯, 1)^T^. **b.** We randomly set the initial abundance *x*_0_ of species at a distance *d* = 0.4 from the desired state. Without control, the state of the microbial community does not reach the desired state *x*_d_. **c. and d.** For the same community and initial abundance as in panel b, we apply the control input generated by the linear MPC (panel c) to the six driver species we identified. This control strategy successfully drives the state of the community to the desired state (panel d). **e., f. and h.** Mean success rate of our control framework as a function of the distance *d* of the initial state from the desired state. Error bars denote the standard error of the mean. Here, the simulation parameters are: *c* = 0.025, *θ*_max_ = 0.05 for panel e, *σ* = 0.8, *θ*_max_ = 0.05 for panel f, and *c* = 0.025, *σ* = 0.8, *θ*_max_ = 0.05 for panel h. **g.** Success rate of our control framework for different proportions of driver species *M*/*N*. Black dots show the success rate of 7700 random communities plotted as a function of the proportion of driver species. Pink shows the mean success rate as a function of the proportion of driver species.

To quantify the success of our control framework on a particular community, we generate 300 initial species abundances that are uniformly distributed at a distance *d* > 0 from the desired state (distance is measured using the Euclidean norm). Then, the *success rate* of our control framework at distance *d* is defined as the proportion of those initial conditions that are driven to the desired state only when the linear MPC is applied to a minimal set of driver species of the community (Fig. 4b-d). Namely, the success rate discards all initial conditions that naturally evolve to the desired state. Finally, we calculated the *mean success rate* by averaging the success rate over 100 randomly constructed ecological networks (see items 7 and 8 of Supplementary Note 8 for details).

The mean success rate of our control framework changes with the distance to the desired state, being close to 1 for small distances regardless of the parameters of the microbial community (Fig. 4e-f). This result agrees well with the theoretical prediction that success is guaranteed provided that the distance to the desired state is small enough. We next investigated how the success rate changes with the distance *d* for different interspecies interaction strengths, and for different connectivities of the ecological network underlying the community. The success rate decreases as the interspecies interaction strength increase, especially for large distances (Fig. 4e). Since increasing the interspecies interaction strength damages the stability of the population dynamics^44^, this result suggests that microbial communities become “harder” to control as they lose stability. The success rate of our control framework is also higher in microbial communities whose ecological networks have lower connectivity (Fig. 4f). Note that, in general, the size of a minimal set of driver species decreases as the network connectivity increases. Therefore, this observations suggest that the success rate may increase as the number of driver species increases. Indeed, regardless of the distance to the desired state, we find that our control framework attains a success rate > 0.8 provided that we drive at least 6 of the 100 species (Fig. 4g). This last result also suggests that the success rate of our control framework can be enhanced by directly controlling a few additional species.

Finally, we investigated the robustness of our control framework to errors in the ecological network used for both identifying the driver species, and for calculating the linear MPC. Note that, despite structural accessibility is insensitive to missing interactions in the ecological network, the calculated linear MPC is not. Additionally, structural accessibility can be lost if some ecological interactions do not really exist in the ecological network. To introduce errors in the ecological network, we randomly rewire each of its edges with probability *p* ∈ [0, 1]. This rewiring probability determines the percentage of error introduced to the ecological network (e.g., *p* = 0.05 corresponds to a 5% error). Our control framework is robust to these errors, in the sense that the success rate deteriorates but remains larger than zero despite large errors (Fig. 4h). However, just a 5% error decreases the success rate in about 30%. This result illustrates that our framework is feasible for controlling large microbial communities provided we have an accurate map of their ecological networks.

## APPLICATION

Mapping the ecological network of a microbial community allow us to identify its driver species. We identified a minimal set of driver species in the gut microbiota of germ-free mice that are pre-colonized with a mixture of human commensal bacterial type strains and then infected with *Clostridium difficile* spores^22^. We identified a minimal set of five driver species in this 14-species community: *R. obeum* (x_1_), *R. mirabilis* (x_12_), *B. ovalus* (x_2_), *C. ramnosum* (x_6_) and *A. muciniphila* (x_10_), see Fig. 5a. We also used the ecological network underlying the core microbiota of the sea sponge *Ircinia oros*^23^, finding ten driver species in this twenty-species community (Fig. 5b).

**Figure 5.**
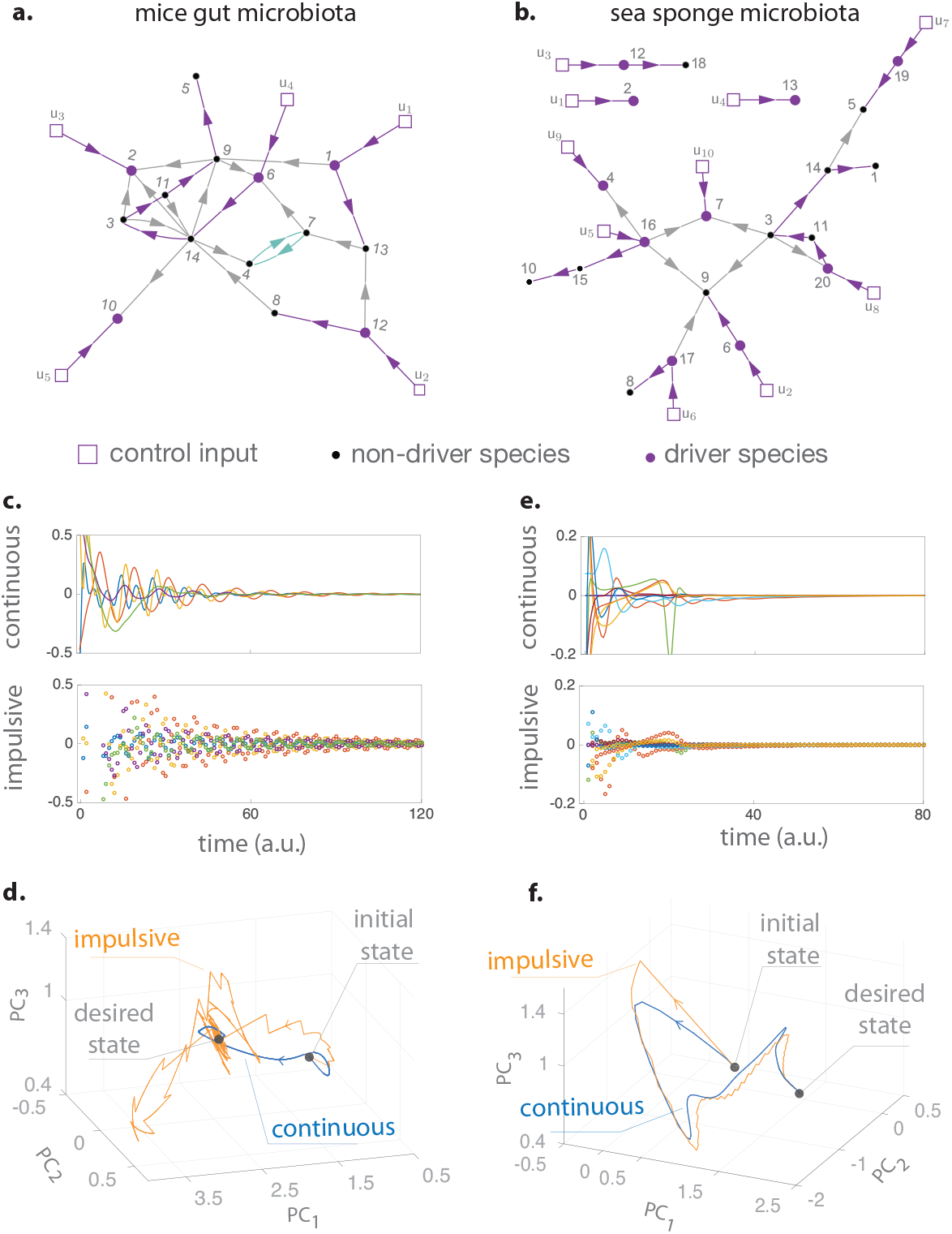
Controlling host-associated microbial communities. **a.** Inferred ecological network of the gut microbiota of germ-free mice pre-colonized with a mixture of human commensal bacterial type strains and then infected with *C. difficile* (species 7). **b.** Inferred ecological network of the core microbiota of the sea sponge *Ircinia oros*. In both networks, self-loops are omitted to improve readability. A minimal set of driver species is shown, providing a disjoint union of paths (purple) and cycles (green) covering all species nodes. Refer to Table 1 in Supplementary Note 7 for the species name. The controlled population dynamics of both microbial communities were simulated using the cGLV equations (see Supplementary Note 7 for details). The intrinsic growth rates were adjusted such that the community has an initial “diseased” equilibrium state *x*_0_ in which one species (*C. difficile* for the mice gut microbiota) is overabundant compared to the rest of species. We chose the desired state *x*_d_ as another equilibrium with a more balanced abundance profile. **c, e.** Control actions obtained using the linear MPC for the impulsive and continuous control schemes. **d, f.** Projection of the high-dimensional abundance profiles (states of the microbial communities) into their first three principal components (PCs). See Supplementary Fig.S1 for the temporal response of each species. The calculated control strategies applied to the driver species succeed in driving the community to the desired state, using either continuous or impulsive control.

We studied by simulation the efficacy of the identified driver species and the linear MPC method for these two microbial communities, assuming that their dynamics are uncertain (see Supplementary Note 7 for details of the dynamics used for the simulation). For the mice gut microbiota, our framework succeeds in driving the whole community from an initial state where *Clostridium difficile* is overabundant, towards a desired state with a better balance of species (Figs. 5c and 5d). Similar results were obtained for controlling the core microbiota of *Ircinia oros*, using the ten identified driver species to drive the twenty species constituting this microbial community (Figs. 5e and 5f). The success of our control framework shows again that the linear MPC method is robust enough to drive microbial communities despite the presence of the perturbations (*w_x_*, *w_u_*).

## DISCUSSION

An influential method to understand and manage complex ecosystems has been identifying species with a “big impact” on the entire ecosystem, leading to notions such as keystone^45, 46^ or core^47^ species. In general, the keystone or core species of an ecosystem are not necessarily its driver species. For example, the driver species of an ecosystem do not depend on their abundance, while the definition of keystone species does depend on the abundance —namely, species whose removal cause a disproportionate deleterious effect relative to their abundance^45^.

It was suggested that notion of *controllability* —the ability to drive a system between any two states— could help predicting the success of ecosystem management strategies^48^. For microbial communities and many other biological systems, it is inadequate to use the notion of controllability because there are states that those systems cannot reach by their nature (e.g., those states corresponding to negative abundances). Additionally, since dynamic models for microbial communities and other complex ecosystems are nonlinear, uncertain, and often very difficult to infer, it is impossible to even test if those systems are controllable or not. The notion of structural accessibility at the basis of our framework overcomes these two limitations, generalizing the control-theoretic notion of accessibility^32^ to systems with uncertain dynamics and impulsive control inputs. As result, our framework allows efficiently controlling microbial communities only knowing their underlying ecological networks. We note that our framework can be used to identify minimal sets of “driver variables” for biological systems beyond microbial communities when their underlying networks are known. For this, we just need to choose the adequate base model^49^ for each class of system. For example, we identified a single “driver protein” in the repressilator^50^ —a synthetic three-gene regulatory network that generates sustained oscillations— allowing us to eliminate those oscillations (Supplementary Note 8 and Fig. S2).

In this paper, we used a maximum matching based algorithm to identify a minimum set of driver species from the ecological network of a given microbial community. In principle, there could be multiple maximum matchings associated with the same network, rendering potentially different minimum sets of driver species. Note that those minimum driver species sets share the same cardinality. We claim that a minimum set of driver species is optimal only in the sense that its cardinality is minimal. If the cost of choosing any species as a driver species is known, one can develop a combinatorial optimization scheme to further pick up the best driver species set. But we feel this is beyond the scope of the current work and hence leave it for future work.

Rather counterintuitively, our mathematical formalism shows that increasing the complexity of the community’s population dynamics (measured by the size of the deformation) can only reduce the number of necessary driver species. In practice, however, increasing the complexity of the dynamics could render the design of the control strategies more difficult. Note that, in general, it can be expected that the design of control strategies becomes more difficult as the number of used driver species decreases (see Remark 9 in Supplementary Note 5). Additionally, we note that despite the minimal number of driver species decreases as the ecological network becomes denser, this condition is only sufficient. Indeed, the minimal number of driver species of a microbial community should be mainly determined by the degree distribution of the ecological network, since the maximum matching size of a directed network is largely determined by its degree distribution^51^.

For large communities with uncertain controlled population dynamics, we calculated the control actions using a linear prediction model with an infinite horizon. More sophisticated control algorithms, such as those based on reinforcement learning^52^ (RL), could provide better performance. Note that RL algorithms typically require specifying a-priori the “driver variables” they can actuate^53^. Our characterization of minimal sets of driver species should help to efficiently apply RL methods for controlling microbial communities and other biological systems. In practice, the performance of the control algorithms can also be improved by using more detailed models that incorporate the dynamics of the susceptibility of species to the control actions (e.g., the pharma cokinetics of prebiotics). In such case, different control actions could be modeled by different pairs {*f*, *g*} in Eqs. (2) or (3), making the conditions for the absence of autonomous elements different for continuous and impulsive control actions. We note that altering the ecological network of a microbial community or obtaining a “simplified” network, in the spirit of Refs.^54^ and ^55^, respectively, could be an alternative and complementary approach to controlling microbial communities (e.g., to reduce the number of necessary driver species).

Note also that in our deterministic framework we don’t consider the effects of stochasticity due to, e.g., immigration in microbial communities. From a theoretical viewpoint, incorporating stochastic effects into the model will turn Eqs. (2) and (3) into controlled stochastic differential equations, which are the material of a different scientific area. To the best of our knowledge, the characterization of the accessibility properties of those class of equations remains an open problem and their analysis become intractable in practice. Indeed, the very notion of an autonomous element —the basis for the concept of accessibility— would need to be reformulated. We consider this is beyond the scope of the current work and call for research activities of the control theory community in this area.

In conclusion, by identifying driver species, our framework shows that an accurate map of the ecological network underlying a microbial community opens the door for an efficient and systematic control. The driver species can be identified despite missing interactions in the ecological network, but our methods to calculate the adequate control actions can be sensitive to them. The design of controllers that are robust to missing interactions will be a necessary step for controlling real microbial communities. To fully harvest the potential benefits of controlling microbial communities a stronger synergy between microbiology, ecology, and control theory will be necessary.

### Data availability

All the experimental datasets analyzed in this study are publicly available.

### Code availability

A Julia implementation of the algorithm for identifying a minimal set of driver species, as well as all other functions necessary to reproduce the results of the paper, is provided at the GitHub repository: https://github.com/mtangulo/DriverSpecies.

